# Functional and comparative genomic analysis of integrated prophage-like sequences in *Candidatus Liberibacter asiaticus*

**DOI:** 10.1101/661967

**Authors:** Marian Dominguez-Mirazo, Rong Jin, Joshua S. Weitz

## Abstract

Huanglongbing (HLB; yellow shoot disease) is a severe worldwide infectious disease for citrus family plants. The pathogen *Candidatus Liberibacter asiaticus* (CLas) is an alphaproteobacterium of the *Rhizobiaceae* family that has been identified as the cause. The virulence of CLas has been attributed, in part, to prophage encoded genes. Prophage and prophage like elements have been identified in 12 of the 15 CLas available genomes, and are classified into three prophage types. Here, we re-examined all 15 CLas genomes using a *de novo* prediction approach and expanded the number of prophage like elements from 16 to 33. Further, we find that all CLas contain at least one prophage-like sequence. Comparative analysis reveals a prevalent, albeit previously unknown, prophage-like sequence type that is a remnant of an integrated prophage. Notably, this remnant prophage is found in the Ishi-1 CLas strain that had previously been reported as lacking prophages. Our findings provide both a resource and new insights into the evolutionary relationship between phage and CLas pathogenicity.

## 1 Introduction

*Candidatus Liberibacter asiaticus* (CLas), has been identified as one of the three Candidatus Liberibacter species that cause HLB disease (HLB: yellow shoot disease, also known as citrus greening disease). HLB is a major threat to the worldwide citrus growing industry [1]. Symptoms of infected trees include leaf mottling, deformed/discolored fruits, premature fruit drop, and premature mortality [2]. It has been suggested that CLas encoded proteins have an inhibitory effect in plant defenses [3, 4], but a comprehensive mechanism of infection is still lacking. No effective disease management practice is currently available. Hence, understanding the genomic composition of CLas and its infection mechanisms is likely to help develop strategies to manage the disease.

As CLas has not been cultivated *in vitro*, its biological study has been based on analyses of DNA extracted from infected plants or insects. To date, a total of 15 genome assemblies are available at the NCBI Genome database. The CLas genome size is ~1.2 Mb with a G+C content of 36.5%; both small genome size and low G+C content are consistent with patterns of obligate intracellular bacteria [5]. The CLas genome includes genes involved in cell motility and active transport [5, 6], despite the presence of these genes, only passive movement has been observed in the phloem sap [6]. Furthermore, prophage sequences have also been identified in multiple CLas strains[7, 8, 9, 10, 11, 12, 13, 14, 15]. The interest in Clas prophage stems from observations that a prophage-encoded peroxidase is an effector that suppresses plant defenses [3]. However, strains reported to lack prophages still induce HLB symptoms [16, 8]. Hence, it has been hypothesized that prophages might contribute to bacterial virulence but are not required for CLas pathogenicity [6].

Prophage regions in CLas genomes are highly variable relative to the rest of the genome [16, 8]. Comparative analyses of prophage sequences suggest endemism in CLas strains [17, 14]. Based on currently available sequence data, CLas prophages have been classified as uniquely belonging to one of three Types, i.e. Type 1, 2 or 3. Current methods to identify prophage in CLas genomes rely on read mapping against known prophages [12, 9]. As a consequence, failure of read-mapping has been interpreted to mean that a genome is prophage free (e.g., as in the Ishi-1 strain [16]). However, a recent study identified the first Type 3 prophage by using BLAST searches with phages SC1 and SC2 as query to locate phage containing contigs, within a CLas genome with incomplete read-mapping to Type 1 or Type 2 prophage [13]. More generally, we are unaware of any systematic analysis of prophage content in CLas genomes using *de novo* prediction approaches.

In this study, we examined the genome sequences of 15 CLas strains and extracted phage sequences using *de novo* prediction tools, including Virsorter [18] and PHASTER [19]. We identified several potential prophage elements not available in databases, of which 5 belong to either Type 1, 2 or 3 CLas phage elements that had not been previously reported. We also identified 12 sequences belonging to a group present in all strains that does not resemble any previously identified phage type. Several analyses show that these sequences do not match Type 1, 2 or 3 prophages. We argue that, based on composition and evolutionary analysis, it is likely that these sequences belong to a different CLas prophage-like sequence, which we term “Type 4”. Multiple lines of evidence suggest that Type 4 prophage-like sequences are remnants of an integrated prophage. The study expands the number of prophage-like elements from 16 to 33. The results will provide a genomic resource for future investigations in the role of prophage in shaping the pathogenicity, ecology, and evolution of CLas.

## 2 Methods

### 2.1 Genome sequences

A total of 15 genome sequences from different CLas strains were obtained from the National Center for Biotechnology Information (NCBI) Nucleotide Database, under the genome ID 1750 (Table 1)(https://www.ncbi.nlm.nih.gov/genome/genomes/1750). All CLas prophages previously reported were also retrieved from NCBI (Table 2).

**Table 1:** *Candidatus Liberibacter asiaticus* genome information. Characteristics of the 15 publicly available CLas genomes: strain, geographic origin, GeneBank Accesion ID, contig number, sequence length, G+C content and, associated phage types. Phage types with asterisks belong to phages with publicly available genomes (See Table 2)

| Strain | Source | Accession ID | Contigs | Length (bp) | GC (%) | Prophage Type |
| --- | --- | --- | --- | --- | --- | --- |
| A4 | Guangdong, China | GCA.000590865.2 | 1 | 1,233,514 | 36.4 | 2 |
| AHCA1 | California, USA | GCA.003143875.1 | 1 | 1,233,755 | 36.63 | 1 |
| FL17 | Florida, USA | GCA.000820625.1 | 3 | 1,227,253 | 36.47 | 1 |
| gxpsy | Guangxi, China | GCA.000346595.1 | 1 | 1,268,237 | 36.57 | 1, 2 |
| HHCA1 | California, USA | GCA.000724755.2 | 239 | 1,150,620 | 36.55 | 2* |
| Ishi-1 | Okinawa, Japan | GCA.000829355.1 | 1 | 1,190,853 | 36.32 | - |
| JXGC | Jiangxi, China | GCA.002216815.1 | 1 | 1,225,162 | 36.40 | 3* |
| psy62 | Florida, USA | GCA.000023765.2 | 1 | 1,227,328 | 36.47 | 1, 2* |
| SGCA1 | California, USA | GCA.003149415.1 | 606 | 233,414 | 36.25 | 1 |
| SGCA5 | California, USA | GCA.001430705.1 | 56 | 1,201,385 | 36.36 | 1* |
| SGpsy | California, USA | GCA.003336865.1 | 1,402 | 769,888 | 36.32 | 1 |
| TX1712 | Texas, USA | GCA.003160765.1 | 48 | 1,203,333 | 36.36 | - |
| TX2351 | Texas, USA | GCA.001969535.1 | 71 | 1,252,002 | 36.52 | - |
| YCpsy | Guangdong, China | GCA.001296945.1 | 9 | 1,233,647 | 36.48 | 1, 3 |
| YNJS7C | Yunnan, China | GCA.003615235.1 | 3 | 1,258,986 | 36.58 | 2, 3 |

**Table 2:**
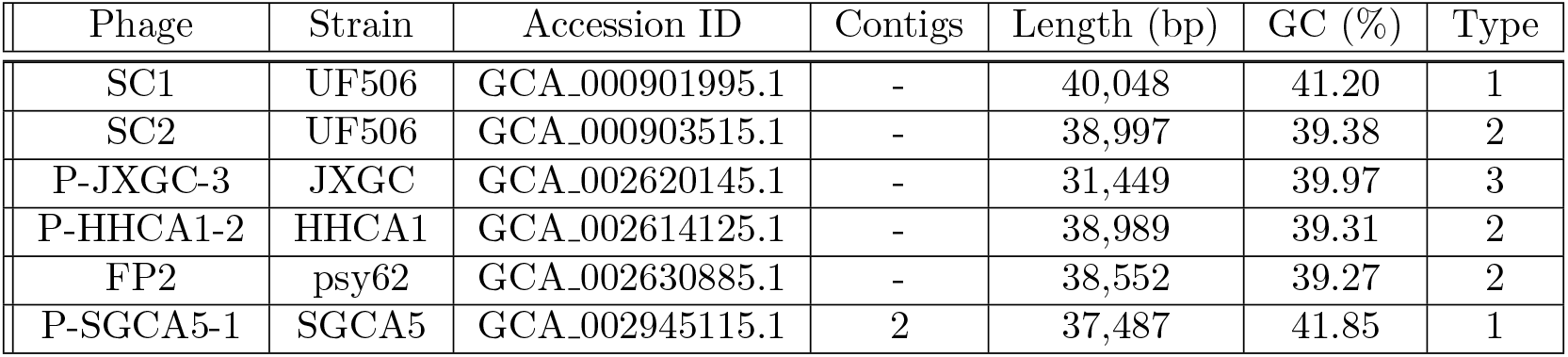
CLas phages genome information. Characteristics of the 5 publicly available genomes of CLas phages: Phage name, associated CLas strain, GeneBank Accesion ID, contig number, sequence length, GC content, and phage type. Empty spaces in the contig column correspond to closed genomes.

### 2.2 Prophage-like sequence identification

All CLas genomes were examined for prophage sequences using the phage identification tools Virsorter[18] and PHASTER[19]. The genomes were imported into Virsorter [18] in the Discovery Environment (CyVerse) and evaluated against the available Refseq database. As PHASTER only works for sequences larger than 1,500 bp, contigs with a size smaller than the required were discarded. Only prophage-like sequences predicted by both tools were kept for further analysis.

### 2.3 Phage classification

Classification of putative prophages began with a comparison to NCBI available phage genomes. Predicted prophages that matched publicly available phage were not subject to further classification. In all other cases, SC1, SC2, and P-JXGC-3 phages were used as representative phages of Type 1, Type 2 and Type 3. Two way nucleotide BLASTs were performed with default parameters with the predicted sequence as query or target. Coverage and Identity values were divided in high, medium or low. Identity (I) was considered high with I ≥ 95, medium with 95 > I ≥ 85, and low with I ¡ 85. Two sets of thresholds were used to evaluate the coverage. When the query sequence was shorter than the target, the Coverage (C) was considered high with C ≥ 90, medium with 90 > C ≥ 75, and low with C < 75. When the query was longer than the target, the Coverage was considered high with C ≥ 33, medium with 33 > C ≥ 25, and low with C < 25. A sequence was classified as a particular phage type if none of its BLAST values was “low” and if it had the highest fraction of “high” scored values. If more than one phage type full-filed the aforementioned conditions, the assigned phage type was the one with the highest identity. The putative prophages that were not classified as Type 1, 2 or 3 were aligned against each other and the formed groups were classified as a single Type 4 group.

### 2.4 CDS prediction, sequence clusterization and gene annotation

For sequence annotation, open reading frames (ORFs) were identified with GeneMarkS-2 [20]. Homology between predicted proteins was obtained using the get_homologues software [21] with default parameters of the Bidirectional Best Hit. The presence-absence analysis of the pangenome was done through a hierarchical cluster in R to group prophage-like sequences according to their shared CDS. Default parameters for the heatmap.2 function from the gplot package (version 3.0.1.1) were used, including “average” as linkage method. The protein predictions were annotated with BLAST against the nr database. The Tandem Repeats Finder [22] was used to find potential attachment sites 5,000 bp upstream and 5,000 bp downstream the predicted prophage sequences.

### 2.5 Code availability

R (v3.3) and Bash were used to generate all figures. Code is available at https://github.com/WeitzGroup/CLas_prophages with https://doi.org/10.5281/zenodo.3238703.

## 3 Results

### 3.1 Identification of novel prophage-like sequences

A total of 35 putative prophage-like sequences were identified among the 15 CLas genomes by Virsorter and PHASTER. At least one sequence was predicted for each of the strains studied (see Table 3) excepting SGCA1 and SGpsy, which may be due to the high fragmentation of their assemblies (see Table 1). Predicted sequences vary highly in length with the shortest sequence having ~2,000 bp and the longest over 65,000 bp. The G+C content of prophage-like sequences can be divided into two groups, those around 40% (similar to the G+C content of Type 1, 2 and 3 prophages), and those around 36% (similar to the CLas value). We compared the predicted sequences to NCBI available phage genomes. Of the 35 predicted sequences, 6 belonged to reported phage genomes. With 3 prophage sequences of HHCA1, 1 of JXGC, 1 of psy62, and 1 of SGCA5, belonging to phages P-HHCA1-2, P-JXGC-3, FP2, and P-SGCA5-1, respectively. There are 29 sequences that are not part of a publicly available prophage genome.

**Table 3:** Putative prophage sequences characteristics. Characteristics of the sequences predicted by Virsorter and PHASTER: CLas strain, sequence ID, contig in which the sequence was found, position in contig, sequence length, and G+C content. Empty values (marked as “−”) in the Contig column indicate the corresponding genome is fully assembled.

| Strain | Sequence ID | Contig | Position | Length (bp) | G+C content (%) |
| --- | --- | --- | --- | --- | --- |
| A4 | A4-a | - | 1,130,506 - 1,149,463 | 18,958 | 35.31 |
| A4 | A4-b | - | 1,189,717 - 1,232,852 | 43,136 | 39.52 |
| AHCA1 | AHCA1-a | - | 1,135,796 - 1,196,595 | 60,800 | 36.66 |
| FL17 | FL17-a | 1 | 1,132,100 - 1,197,885 | 65,786 | 36.54 |
| FL17 | FL17-b | 2 | 109 - 15,184 | 15,076 | 43.39 |
| FL17 | FL17-c | 3 | 76 - 13,493 | 13,418 | 39.75 |
| gxpsy | gxpsy-a | - | 1,118,915 - 1,137,878 | 18,964 | 35.36 |
| gxpsy | gxpsy-b | - | 1,178,134 - 1,226,132 | 47,999 | 39.28 |
| gxpsy | gxpsy-c | - | 1,226,133 - 1,266,293 | 40,161 | 41.50 |
| HHCA1 | HHCA1-a | 217 | 189 - 6,282 | 6,094 | 37.12 |
| HHCA1 | HHCA1-b | 229 | 160 - 8,846 | 8,687 | 41.56 |
| HHCA1 | HHCA1-c | 231 | 11 - 13,045 | 13,035 | 40.84 |
| HHCA1 | HHCA1-d | 239 | 11,351 - 39,707 | 28,357 | 39.26 |
| Ishi-1 | Ishi-1-a | - | 1,127,556 - 1,144,572 | 17,017 | 35.31 |
| JXGC | JXGC-a | - | 1,130,487 - 1,149,448 | 18,962 | 35.35 |
| JXGC | JXGC-b | - | 1,189,703 - 1,224,971 | 35,269 | 40.20 |
| psy62 | psy62-a | - | 948 - 11,549 | 10,602 | 40.40 |
| psy62 | psy62-b | - | 1,180,444 - 1,226,282 | 45,839 | 40.10 |
| psy62 | psy62-c | - | 1,132,029 - 1,149,043 | 17,015 | 35.30 |
| SGCA5 | SGCA5-a | 22 | 802 - 4,833 | 4,032 | 37.03 |
| SGCA5 | SGCA5-b | 43 | 3,028 - 17,448 | 14,421 | 44.39 |
| TX1712 | TX1712-a | 3 | 51,093 - 68,108 | 17,016 | 35.30 |
| TX1712 | TX1712-b | 17 | 127 - 27,513 | 27,387 | 41.80 |
| TX1712 | TX1712-c | 35 | 1,026 - 7,842 | 6,817 | 41.94 |
| TX2351 | TX2351-a | 66 | 677 - 10,598 | 9,922 | 41.14 |
| TX2351 | TX2351-b | 67 | 438 - 11,367 | 10,930 | 39.11 |
| TX2351 | TX2351-c | 69 | 1,515 - 7,899 | 6,385 | 37.79 |
| TX2351 | TX2351-d | 70 | 288 - 13,018 | 12,731 | 40.60 |
| YCPsy | YCPsy-a | 2 | 692,231 - 711,192 | 18,962 | 35.35 |
| YCPsy | YCPsy-b | 3 | 13 - 21,283 | 21,271 | 43.00 |
| YCPsy | YCPsy-c | 6 | 30 - 2,207 | 2,178 | 39.99 |
| YCPsy | YCPsy-d | 7 | 1,952 - 7,990 | 6,039 | 41.91 |
| YNJS7C | YNJS7C-a | 1 | 1,125,779 - 1,142,794 | 17,016 | 35.31 |
| YNJS7C | YNJS7C-b | 2 | 735 - 39,556 | 38,822 | 41.79 |
| YNJS7C | YNJS7C-c | 3 | 12,161 - 31,475 | 19,315 | 39.81 |

### 3.2 Classification of prophage-like predicted sequences

Of the 29 predicted phage sequences, a total of 13 were successfully classified into Type 1, 2, or 3 (see Methods, Table 4). As previously described, strains psy62, gxpsy, FL17, and YCPsy have Type 1 classified sequences. Strains A4 and gxpsy have a Type 2 classified sequence. Strain YNJS7C harbours a sequence that resembles a Type 3 prophage [12, 23]. Novel prophage-like sequences of Type 1 were found in strains TX1712 and TX2351, while Type 2 sequences were also found in strain TX2351. Contrary to a previous report, no YNJS7C sequences were classified as Type 2 [15], but instead a Type 1 sequence was identified. Although they seemed to resemble the representative phages, 4 sequences remained unclassified due to their small size. Finally, 12 sequences had similar characteristics between them, but 10 had no apparent resemblance to the representative phages SC1, SC2, and P-JXGC-3, while a small subsequence of the other two (AHCA1-a and FL17-a) partially resembled the representative phages (see Table 4).

**Table 4:** Classification of novel prophage-like sequences into Type 1, 2 and 3. The novel predicted sequences were aligned against representative phages SC1, SC2 and P-JXGC-3 of Type 1, 2 and 3. Coverage (C) values of two-way local alignments are presented on this table. Green, yellow, and red cells stand for high, medium and low alignment values. Note that the threshold values used to classify coverage (C) vary according to sequence length and position in BLAST (either query or target) (see methods). White cells with “NA” values stand for no significant hits. Sequences were divided in either Type 1 (red), Type 2 (blue), Type 3 (green), unclassified sequences with resemblance to the representative phages (black), or unclassified without resemblance (gray).

|  | Type 1 |  |  |  | Type 2 |  |  |  | Type 3 |  |  |  |
| --- | --- | --- | --- | --- | --- | --- | --- | --- | --- | --- | --- | --- |
|  | SC1 |  |  |  | SC2 |  |  |  | P-JXGC-3 |  |  |  |
|  | Target |  | Query |  | Target |  | Query |  | Target |  | Query |  |
| Value (%) | C | I | C | I | C | I | C | I | C | I | C | I |
| SC1 | 100 | 100.0 | 100 | 100.0 | 52 | 98.49 | 52 | 98.49 | 42 | 97.89 | 53 | 97.89 |
| SC2 | 52 | 98.49 | 52 | 98.49 | 100 | 100.0 | 100 | 100.0 | 47 | 97.43 | 58 | 97.43 |
| P-JXGC-3 | 53 | 97.89 | 42 | 97.89 | 58 | 97.43 | 47 | 97.43 | 100 | 100.0 | 100 | 100.0 |
| FL17-b | 97 | 96.29 | 37 | 96.29 | 4 | 96.11 | 1 | 96.11 | NA | NA | NA | NA |
| FL17-c | 98 | 98.62 | 34 | 98.62 | 62 | 96.35 | 22 | 96.35 | 63 | 98.15 | 27 | 98.15 |
| gxpsy-c | 89 | 97.83 | 92 | 97.83 | 44 | 96.38 | 45 | 96.38 | 41 | 98.47 | 52 | 98.47 |
| psy62-b | 71 | 99.80 | 81 | 99.80 | 29 | 98.07 | 33 | 98.07 | 19 | 98.80 | 28 | 98.80 |
| TX1712-b | 100 | 99.81 | 72 | 99.80 | 34 | 98.19 | 27 | 98.19 | 22 | 99.08 | 19 | 98.94 |
| TX2351-a | 100 | 99.92 | 25 | 99.92 | 89 | 99.54 | 23 | 99.54 | 89 | 98.0 | 28 | 98.0 |
| YCPsy-b | 93 | 97.88 | 52 | 97.88 | 17 | 99.63 | 11 | 99.63 | 8 | 99.94 | 5 | 99.94 |
| YNJS7C-b | 89 | 97.86 | 87 | 97.86 | 47 | 97.26 | 45 | 97.26 | 43 | 96.29 | 52 | 96.29 |
| A4-b | 46 | 96.01 | 48 | 96.01 | 90 | 98.28 | 98 | 98.28 | 43 | 94.99 | 58 | 95.0 |
| gxpsy-b | 50 | 98.22 | 53 | 98.22 | 87 | 98.64 | 97 | 98.64 | 44 | 99.49 | 61 | 99.49 |
| TX2351-b | 8 | 92.55 | 5 | 92.55 | 100 | 99.83 | 28 | 99.83 | NA | NA | NA | NA |
| TX2351-d | 92 | 97.23 | 29 | 97.23 | 100 | 99.91 | 33 | 99.91 | 99 | 95.97 | 40 | 95.97 |
| YNJS7C-c | 85 | 95.91 | 42 | 95.91 | 94 | 96.03 | 47 | 96.03 | 100 | 98.11 | 61 | 98.11 |
| TX1712-c | 100 | 99.31 | 17 | 99.31 | 100 | 99.93 | 18 | 99.93 | 100 | 98.27 | 22 | 98.27 |
| TX2351-c | 79 | 98.07 | 13 | 98.07 | 100 | 99.83 | 17 | 99.83 | 81 | 91.83 | 16 | 91.83 |
| YCPsy-c | 100 | 97.80 | 6 | 97.80 | 100 | 97.80 | 6 | 97.80 | 99 | 99.80 | 7 | 99.80 |
| YCPsy-d | 99 | 97.73 | 15 | 97.73 | 99 | 98.11 | 16 | 98.11 | 99 | 99.42 | 20 | 99.42 |
| AHCA1-a | 2 | 93.87 | 4 | 93.87 | 4 | 96.18 | 6 | 96.18 | 4 | 98.02 | 7 | 98.02 |
| FL17-a | 6 | 95.72 | 13 | 95.72 | 5 | 92.50 | 11 | 92.51 | 3 | 94.84 | 6 | 94.85 |
| A4-a | NA | NA | NA | NA | NA | NA | NA | NA | NA | NA | NA | NA |
| gxpsy-a | NA | NA | NA | NA | NA | NA | NA | NA | NA | NA | NA | NA |
| HHCA1-a | NA | NA | NA | NA | NA | NA | NA | NA | NA | NA | NA | NA |
| ishi1-a | NA | NA | NA | NA | NA | NA | NA | NA | NA | NA | NA | NA |
| JXGC-a | NA | NA | NA | NA | NA | NA | NA | NA | NA | NA | NA | NA |
| psy62-c | NA | NA | NA | NA | NA | NA | NA | NA | NA | NA | NA | NA |
| SGCA5-a | NA | NA | NA | NA | NA | NA | NA | NA | NA | NA | NA | NA |
| TX1712-a | NA | NA | NA | NA | NA | NA | NA | NA | NA | NA | NA | NA |
| YCPsy-a | NA | NA | NA | NA | NA | NA | NA | NA | NA | NA | NA | NA |
| YNJS7C-a | NA | NA | NA | NA | NA | NA | NA | NA | NA | NA | NA | NA |

### 3.3 Identification of a new Type of CLas prophage-like sequences

We next set out to identify the “type” of prophage for the unclassified prophage-like sequences found via our *de novo* approach. Of the 12 unclassified sequences that do not resemble the three known phage types, 10 are highly similar to each other and belong to a candidate Type 4 prophage-like sequence that includes the Ishi-1 prophage prediction (Table 5). Our evidence that these prophage-like sequences belong to a new Type is as follows. First, there is no apparent resemblance to prophages of Type 1, 2, or 3 at the nucleotide level, nor at the amino acid level. Second, the length of the Type 4 sequences is about half the size of representative phages of Type 1, 2 and 3 (see Table 6). Third, in contrast to the previously known prophage types (with G+C content of ~40%), Type 4 sequences have a G+C content of 35%-37%, which is closer to that of the host whose G+C content is ≈36% (see Table1 and Table 6).

**Table 5:**
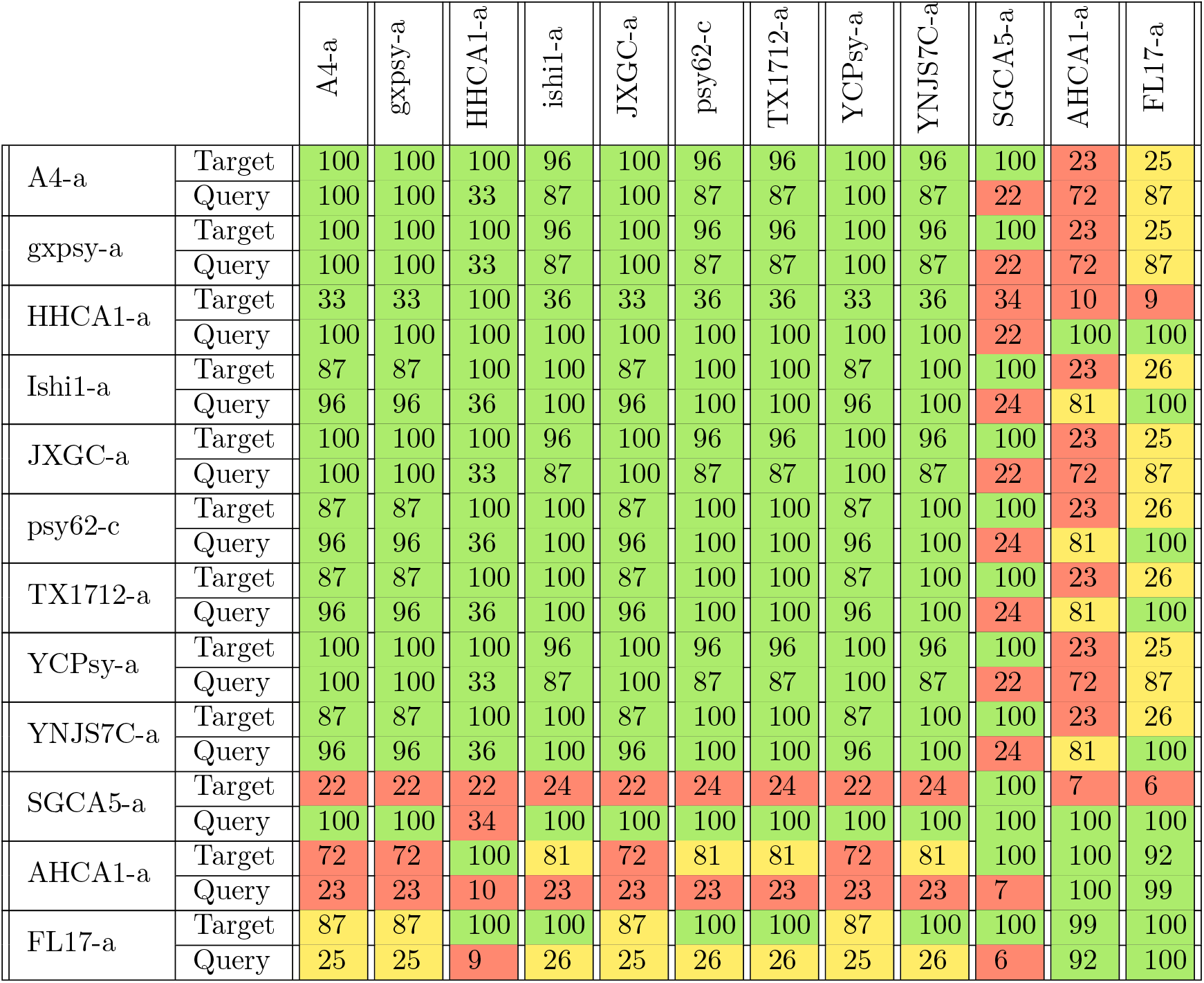
Resemblance between new Type 4 sequences. The 12 predicted sequences that did not resemble any of the representative phages of type 1, 2 and 3 were compared between them. Coverage (C) values of two-way local alignments are presented on this table. Green, yellow, and red cells stand for high, medium and low values, respectively (see methods).

**Table 6:** Putative prophage sequences characteristics and classification. Characteristics of the predicted sequences: Sequence ID, length, G+C content, name (either previously assigned or following the proposed in [12]), and prophage or prophage-like sequence type. Asterik (*) marked sequences AHCA1-a/F1 and AHCA1-a/F2 are different fragments of sequence AHCA1-a. Similarly FL17-a/F1 and FL17-a/F2 are different fragments of sequence FL17-a.

| Sequence | Length (bp) | G+C content (%) | Associated Name | Prophage Type |
| --- | --- | --- | --- | --- |
| FL17-a/F2* | 4,021 | 37.8 | P-FL17-1 | 1 |
| FL17-b | 15,076 | 43.39 | P-FL17-1 | 1 |
| FL17-c | 13,418 | 39.75 | P-FL171-1 | 1 |
| gxpsy-c | 40,161 | 41.50 | P-gxpsy-1 | 1 |
| psy62-b | 45,839 | 40.10 | FP1 / P-psy62-1 | 1 |
| SGCA5-b | 14,421 | 44.39 | P-SGCA5-1 | 1 |
| TX1712-b | 27,387 | 41.80 | P-TX1712-1 | 1 |
| TX2351-a | 9,922 | 41.14 | P-TX2351-1 | 1 |
| YCPsy-b | 21,271 | 43.00 | P-YCPsy-1 | 1 |
| YNJS7C-b | 38,822 | 41.79 | P-YNJS7C-1 | 1 |
| A4-b | 43,136 | 39.52 | P-A4-2 | 2 |
| gxpsy-b | 47,999 | 39.28 | P-gxpy-2 | 2 |
| HHCA1-b | 8,687 | 41.56 | P-HHCA1-2 | 2 |
| HHCA1-c | 13,035 | 40.84 | P-HHCA1-2 | 2 |
| HHCA1-d | 28,357 | 39.26 | P-HHCA1-2 | 2 |
| psy62-a | 10,602 | 40.40 | FP2 / P-psy62-2 | 2 |
| TX2351-b | 10,930 | 39.11 | P-TX2351-2 | 2 |
| TX2351-d | 12,731 | 40.60 | P-TX2351-2 | 2 |
| AHCA1-a/F2* | 2,334 | 39.3 | P-AHCA1-3 | 3 |
| JXGC-b | 35,269 | 40.20 | P-JXGC-3 | 3 |
| YNJS7C-c | 19,315 | 39.81 | P-YNJS7C-3 | 3 |
| A4-a | 18,958 | 35.31 |  | 4 |
| gxpsy-a | 18,964 | 35.36 |  | 4 |
| HHCA1-a | 6,094 | 37.12 |  | 4 |
| Ishi-1-a | 17,017 | 35.31 |  | 4 |
| JXGC-a | 18,962 | 35.35 |  | 4 |
| psy62-c | 17,015 | 35.30 |  | 4 |
| TX1712-a | 17,016 | 35.30 |  | 4 |
| YCPsy-a | 18,962 | 35.35 |  | 4 |
| YNJS7C-a | 17,016 | 35.31 |  | 4 |
| AHCA1-a/F1* | 13,713 | 35.5 |  | 4 |
| FL17-a/F1* | 14,507 | 35.4 |  | 4 |
| SGCA5-a | 4,032 | 37.03 |  | 4 |
| TX1712-c | 6,817 | 41.94 |  | Unknown |
| TX2351-c | 6,385 | 37.79 |  | Unknown |
| YCPsy-c | 2,178 | 39.99 |  | Unknown |
| YCPsy-d | 6,039 | 41.91 |  | Unknown |

The other 2 predicted sequences, FL17-a and AHCA1-a, contain ~80% of the Type 4 sequence (see Table 5) followed by bacterial genes and a fragment highly resembling a part of a Type 1 and a Type 3 prophage genome, respectively (see Figure 1, Table 4). Suppl. Table 1 shows alignment values for the 12 unclassified sequences using the Type 4 region of AHCA1-a and FL17-a.

**Figure 1:**
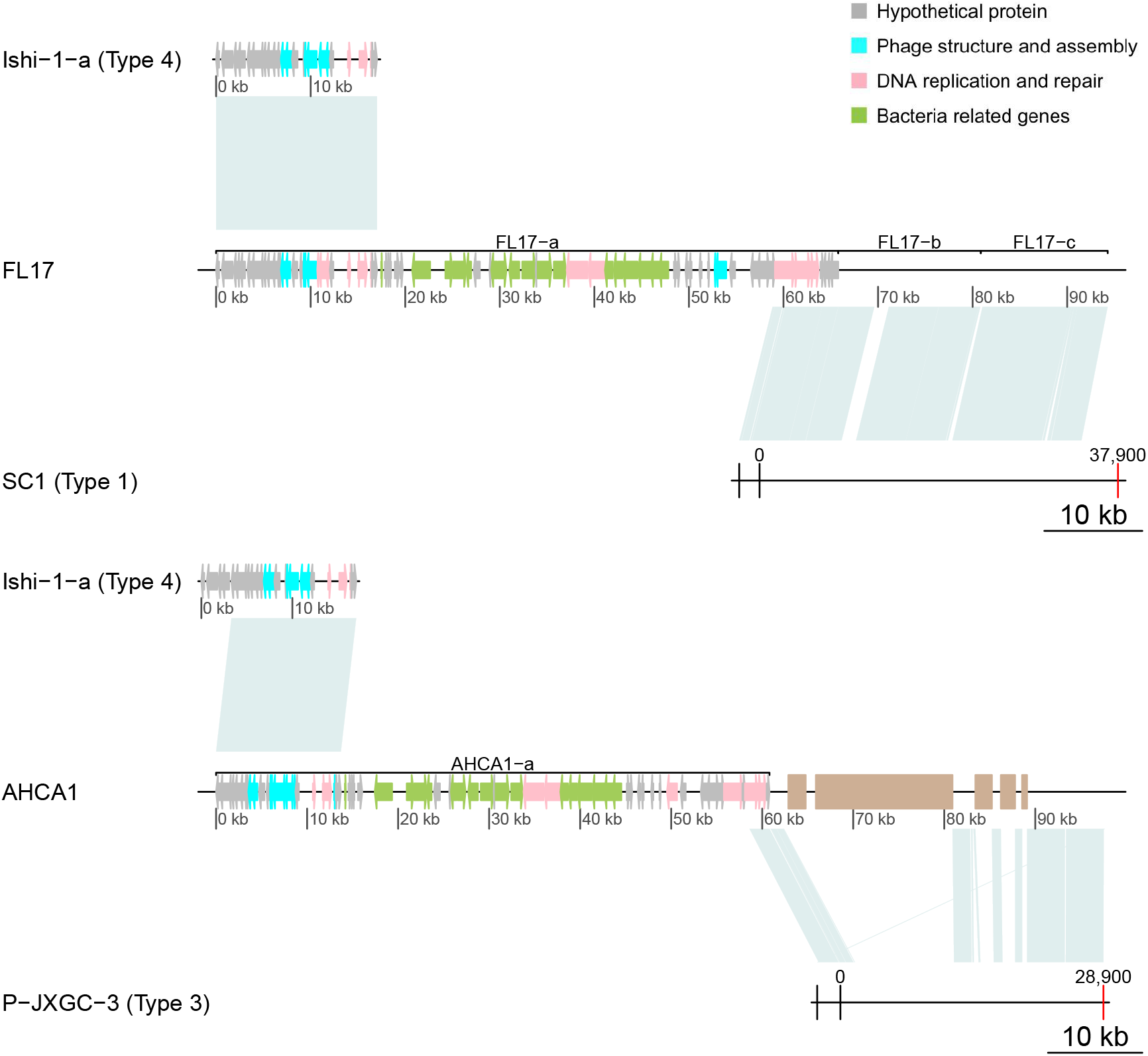
Reconstruction of Type 1 and 3 prophage-like sequences using AHCA1-a, FL17-a and nearby regions. (a) Sequence FL17-a is formed by a Type 4 sequence followed by bacterial genes and a region resembling a Type 1 prophage. Joining sequences FL17-a,b and c reconstructs in its majority a Type 1 prophage sequence with organization similar to the predicted for the integration of SC1 [7]. (b) AHCA-1 contains a Type 4 sequence followed by bacterial genes and a region highly resembling a Type 3 prophage. Using nearby regions in the AHCA1 genome a Type 3 prophage can be reconstructed with an organization similar to the predicted for the integration of P-JXGC-3[13]. Brown rectangles represent assembly gaps.

The prophage fragment at the end of FL17-a (end of contig 1) joined to Type 1 sequences FL17-b (contig 2) and FL17-c (contig 3) reconstructs the sequence of a Type 1 prophage (Figure 1a) with a sequence organization similar to the reported for the chromosomal integration of phage SC1 [7]. Likewise, the last part of the AHCA1-a sequence and the following region found after a genome assembly gap reconstruct a Type 3 prophage with the sequence organization predicted for P-JXGC-3 [13] (Figure 1b). Only a Type 1 prophage has been reported in strain AHCA1 [9], but the sequence organization of the integrated prophage (Figure 1) and the presence of a hypothetical protein (YP_007011137.1) only found in prophages of Type 2 and 3 (see Supl. Table 3), suggest the CLas strain AHCA1 harbours a Type 3 prophage sequence. The truncation of the Type 1 and 3 prophage sequences and their closeness to the Type 4 prophage-like sequence in strains FL17 and AHCA1 explains the incorrect concatenation of these sequences into a single prediction by Virsorter and PHASTER. Table 6 shows all predicted prophage-like sequences with their assigned type and sequence characteristics.

### 3.4 Pan-genome analysis

In order to evaluate the robustness of the classification identified in prior sections, we analyzed the pan-genome content of prophages through a clustering approach (Figure 2). Clustering of gene content in prophage reveals a separation between the 4 types of sequences. All Type 1 sequences cluster together except for TX2351-a. Type 2 and Type 3 sequences appear nearby in the clustering. The Type 2 putative prophage for strain TX2351 clusters with Type 3 sequences. The two mistaken assignments might result from the short size of sequences and the overall close relationship between prophage Types 1, 2, and 3. All sequences of Type 4 cluster together as the most distant group. The clusterization of prophage-like sequences based on gene content predicts relationships that roughly agree with the classification.

**Figure 2:**
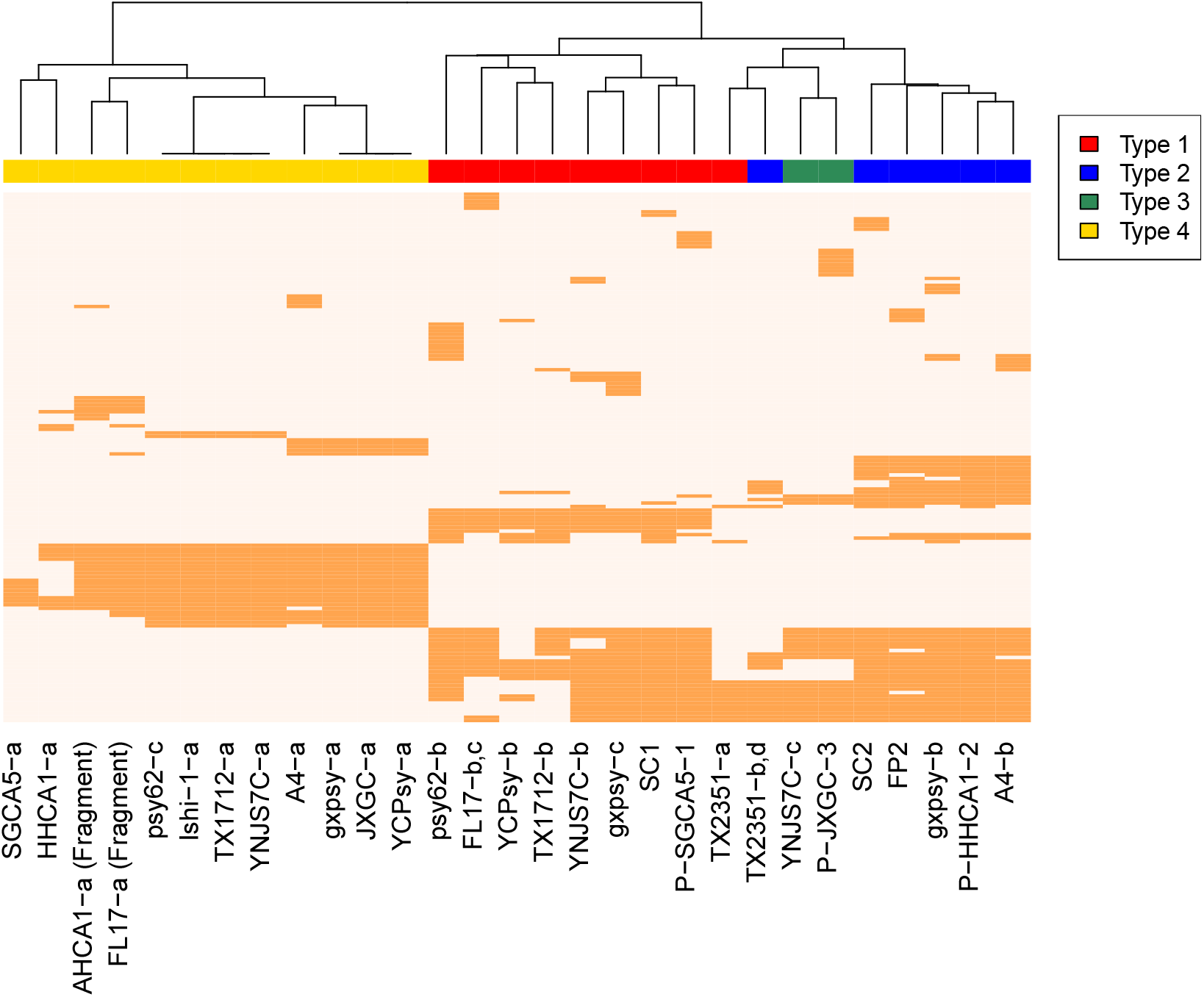
Gene composition in prophage-like sequences. Presence-absence heatmap of predicted pangenome proteins for the prophage-like sequences and database available phages. A clusterization approach was used to obtain relationships between sequences regarding their gene-content. Colors stand for sequence classification, red as Type 1, blue Type 2, green Type 3, and yellow Type 4. Sequences from the same strain that were classified as the same phage type e.g. FL17 type 1, and TX2351 type 2, were used as a single phage for gene content analysis. Unclassified sequences were excluded from the analysis.

Table 7 presents a summary of the associated prophage elements in the 15 CLas strains including the previously reported and the ones introduced in this study. There are 5 newly reported elements that belong to either Type 1, 2, or 3 prophages (see section 3.2), and 12 Type 4 prophage-like sequences. Our findings expand the known prophage elements in CLas from 16 to 33.

**Table 7:** *Candidatus Liberibacter asiaticus* genome information adding new prophagelike sequences. Characteristics of the 15 publicly available CLas genomes: strain, geographic origin, GeneBank Accesion ID, contig number, sequence length, G+C content and, associated prophage types. Prophage types in red belong to types found in this study.

| Strain | Source | Accession ID | Contigs | Length (bp) | GC (%) | Prophage Type |
| --- | --- | --- | --- | --- | --- | --- |
| A4 | Guangdong, China | GCA_000590865.2 | 1 | 1,233,514 | 36.4 | 2, 4 |
| AHCA1 | California, USA | GCA_003143875.1 | 1 | 1,233,755 | 36.63 | 1, 3, 4 |
| FL17 | Florida, USA | GCA_000820625.1 | 3 | 1,227,253 | 36.47 | 1, 4 |
| gxpsy | Guangxi, China | GCA_000346595.1 | 1 | 1,268,237 | 36.57 | 1, 2, 4 |
| HHCA1 | California, USA | GCA_000724755.2 | 239 | 1,150,620 | 36.55 | 2, 4 |
| Ishi-1 | Okinawa, Japan | GCA_000829355.1 | 1 | 1,190,853 | 36.32 | 4 |
| JXGC | Jiangxi, China | GCA_002216815.1 | 1 | 1,225,162 | 36.40 | 3, 4 |
| psy62 | Florida, USA | GCA_000023765.2 | 1 | 1,227,328 | 36.47 | 1, 2, 4 |
| SGCA1 | California, USA | GCA_003149415.1 | 606 | 233,414 | 36.25 | 1 |
| SGCA5 | California, USA | GCA_001430705.1 | 56 | 1,201,385 | 36.36 | 1, 4 |
| SGpsy | California, USA | GCA_003336865.1 | 1,402 | 769,888 | 36.32 | 1 |
| TX1712 | Texas, USA | GCA_003160765.1 | 48 | 1,203,333 | 36.36 | 1,4 |
| TX2351 | Texas, USA | GCA_001969535.1 | 71 | 1,252,002 | 36.52 | 1,2 |
| YCPsy | Guangdong, China | GCA_001296945.1 | 9 | 1,233,647 | 36.48 | 1, 3, 4 |
| YNJS7C | Yunnan, China | GCA_003615235.1 | 3 | 1,258,986 | 36.58 | 1, 2, 3, 4 |

### 3.5 Functional annotation of Type 4 prophage-like sequences

In order to identify characteristic features of Type 4 CLas prophage-like sequences, we functionally annotated the representative prophage-like sequence Ishi-1-a (Figure 3). Annotation of Ishi-1-a revealed 26 putative CDS. Of them, 14 ORFs correspond to hypothetical proteins. Moreover, Ishi-1-a contains several ORFs that present premature stop codons. These ORFs code mainly for fragments of genes with phage structure and assembly functions. Suppl. Table 2 contains the detailed list of the Ishi-1-a sequence annotation.

**Figure 3:**
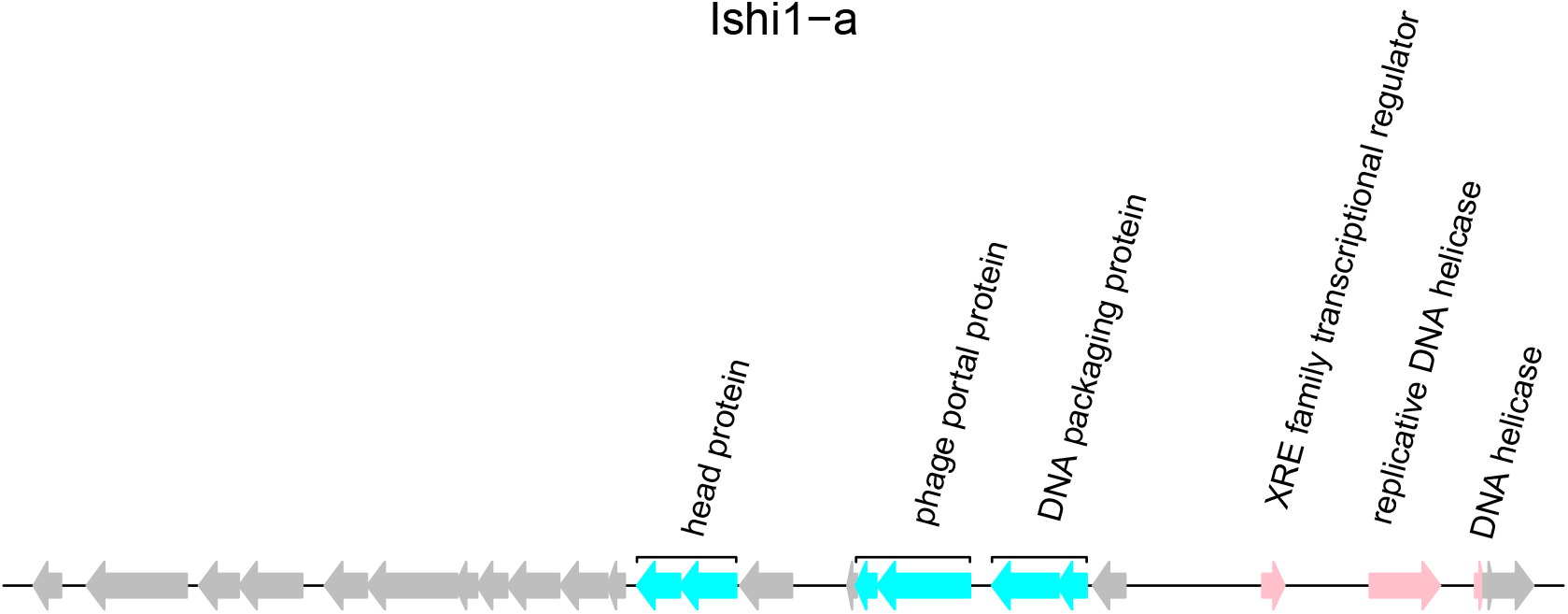
Annotation of representative sequence Ishi-1-a of Type 4. Predicted CDS of Ishi-1-a, with ORFs color coded to gene predicted functions.

In Ishi-1 the sequence of a head protein found in *Candidatus Liberibacter solanacearum* is fragmented into ORF12 and ORF13. Three CDS (ORF15, ORF16, and ORF17) resemble different parts of a portal protein also in *Candidatus Liberibacter solanacearum*. ORF18 and ORF19 are remnants of a DNA packaging protein similar to one present in *Candidatus Liberibacter africanus*. There are three ORFs involved in DNA replication: a gene that codes for a transcriptional regulator of the XRE family (ORF22) and two fragments (ORF23 and ORF24) of an helicase coding gene. Furthermore, no attachment sites were found in the bacterial genome near the Type 4 sequences.

The disruption of viral particle forming genes, along with the absence of att sites, the G+C content closely resembling the one of the bacterial host, and the small sequence size, imply Type 4 sequences are remnants of a prophage integrated into the bacterial genome.

### 3.6 Evolutionary origins of prophage-like sequence Type 4 in CLas

To assess the evolutionary origin of the Type 4 prophage-like sequence, and its time of insertion into the bacterial genome, we searched for the Type 4 sequences in CLas related species. The uncultivated *Candidatus Liberibacter africanus* (CLaf) and *Candidatus Liberibacter americanus* (CLam) species are closely related to CLas and have been identified as less prevalent causative agents of HLB [1]. According to nucleotide and protein alignments, the prophage-like sequence of Type 4 is partially present in both CLaf and CLam (Figure 4). The presence of the Type 4 sequence in these genomes indicated the integration of the Type 4 prophage-like sequence in the Liberibacter genome occurred at least 309 Myr ago (estimated time of lineage divergence for the CLas / CLaf and the CLam clades [24]). The presence of this sequence in all of the HLB known causative agents could suggest a relationship between Type 4 prophage-like sequences and bacterial virulence.

**Figure 4:**
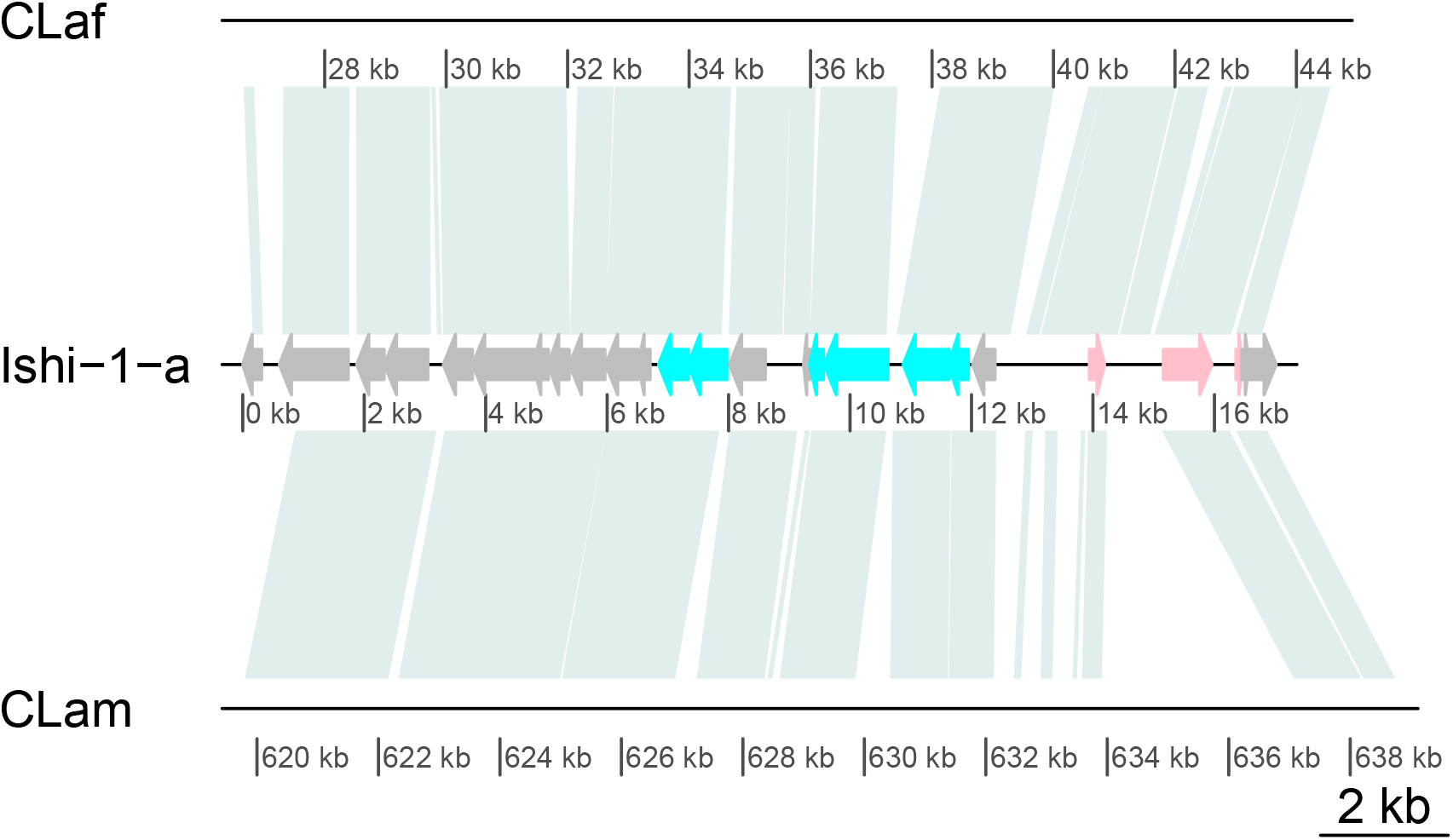
Presence of prophage-like sequence Type 4 in related Candidatus Liberibacter species. Alignment visualization of the Type 4 prophage-like sequence Ishi-1-a against genomes of *Candidatus Liberibacter africanus* (CLaf, GCA_001021085.1) and *Candidatus Liberibacter americanus* (CLam, GCA_000496595.1).

## 4 Discussion

SC1 and SC2 phages were first discovered in 2011 [7]. Since then, the release of a new CLas genome usually includes analysis of prophage types, e.g. [15, 11, 10], whether of Type 1, 2, or 3. These phage types can be used to compare evolutionary dynamics [12, 9], and to predict the origin of strains [17, 14, 9]. Nonetheless, prior work has leveraged read-mapping based approaches to identify new prophage-like sequences, albeit those that resemble Type 1 or 2 phage. Indeed, even the discovery of a Type 3 prophage [13] was due to an incomplete mapping of the known phages. This bias has led to categorization of multiple strains as prophage-lacking [16, 8, 13], and to partially reduce the importance of prophage sequences in the pathogenicity of CLas [8].

Here, we utilized a *de novo* based search to identify new prophage sequences in CLas genomes. Our strategy allowed us to confirm the presence of publicly available phage genomes, new prophages of Types 1, 2, and 3, as well as novel prophage-like sequences that did not resemble previously identified phage types. Subsequent classification analysis identified 12 sequences as part of new Type 4 prophage-like sequences that are found in 12 of the 15 CLas strains. Sequence alignment and gene content analyses support the hypothesis that Type 4 is a new sequence unrelated to the previously identified prophages. Several characteristics suggest Type 4 sequences are remnants of a prophage integrated in the bacterial genome. These characteristics include the disruption of viral particle forming genes, sequence size, the G+C content closely resembling the one of the bacterial host, and the failure to find potential att sites. The presence of a Type 4 sequence in almost all evaluated genomes challenges the common view of CLas strains lacking prophage or prophage-like sequences, and further supports the hypothesis of a mechanistic relationship between prophage–whether active or remnants- and bacterial pathogenicity.

In summary, by adopting a *de novo* prediction approach, we have significantly expanded the diversity of prophage-like sequences found in CLas. Moving forward, combining reference-based and *de novo* approaches is likely to contribute to understanding of the diversity, function and evolution of phage on CLas bacteria. As in other comparative studies of phage, many of the functions of genes in prophage and prophage-like sequences remain unknown. Investigation of the functions of the hypothetical proteins present in prophage-like sequences in bacterial genomes might be the key to further understand the mechanisms of CLas virulence and pathogenicity. We hope that the present study of Clas and associated prophage-like sequences will provide a new approach to identify both causes and solutions to Huanglongbing disease.

## 5 Acknowledgements

The authors thank Burton H. Singer for discussions, and Daniel Muratore for code review. We also thank Sarah R. Bordenstein, Brittany A. Leigh, and Simon Roux for feedback on the manuscript.

