## Supplemental Material for "Functional and comparative genomic analysis of integrated prophage-like sequences in *Candidatus Liberibacter asiaticus*"

### 1 Supplemental Material

| | | | A4-a | AHCA1-a (Fragment) | FL17-a (Fragment) | HHCA1-a | JXGC-a | TX1712-a | YCPsy-a | YNJS7C-a | gxpsy-a | Ishi-1-a | psy62-c | SGCA5-a |
|---|---|---|---|---|---|---|---|---|---|---|---|---|---|---|
| A4-a | Target | C | 100 | 100 | 99 | 100 | 100 | 96 | 100 | 96 | 100 | 96 | 96 | 100 |
| | | I | 100 | 99.8 | 99.7 | 99.2 | 99.8 | 99.7 | 99.8 | 99.8 | 99.8 | 99.8 | 99.7 | 99.8 |
| | Query | C | 100 | 72 | 76 | 33 | 100 | 87 | 100 | 87 | 100 | 87 | 87 | 22 |
| | | I | 100 | 99.8 | 99.7 | 99.2 | 99.8 | 99.7 | 99.8 | 99.8 | 99.8 | 99.8 | 99.7 | 99.8 |
| AHCA1-a (Fragment) | Target | C | 72 | 100 | 95 | 100 | 72 | 81 | 72 | 81 | 72 | 81 | 81 | 100 |
| | | I | 99.8 | 100 | 99.9 | 99.6 | 99.9 | 99.9 | 99.9 | 99.9 | 99.9 | 99.9 | 99.9 | 99.9 |
| | Query | C | 100 | 100 | 100 | 45 | 100 | 100 | 100 | 100 | 100 | 100 | 100 | 30 |
| | | I | 99.8 | 100 | 99.9 | 99.6 | 99.9 | 99.9 | 99.9 | 99.9 | 99.9 | 99.9 | 99.9 | 99.9 |
| FL17-a (Fragment) | Target | C | 76 | 100 | 100 | 100 | 76 | 85 | 76 | 85 | 76 | 85 | 85 | 100 |
| | | I | 99.7 | 99.9 | 100 | 99.7 | 100 | 99.9 | 100 | 99.9 | 99.9 | 99.9 | 99.9 | 99.9 |
| | Query | C | 99 | 95 | 100 | 43 | 99 | 99 | 99 | 99 | 99 | 99 | 99 | 28 |
| | | I | 99.7 | 99.9 | 100 | 99.7 | 100 | 99.9 | 100 | 99.9 | 99.9 | 99.9 | 99.9 | 99.9 |
| HHCA1-a | Target | C | 33 | 45 | 43 | 100 | 33 | 36 | 33 | 36 | 33 | 36 | 36 | 34 |
| | | I | 99.2 | 99.6 | 99.7 | 100 | 99.7 | 99.7 | 99.7 | 99.7 | 99.7 | 99.7 | 99.7 | 99 |
| | Query | C | 100 | 100 | 100 | 100 | 100 | 100 | 100 | 100 | 100 | 100 | 100 | 22 |
| | | I | 99.2 | 99.6 | 99.7 | 100 | 99.7 | 99.7 | 99.7 | 99.7 | 99.7 | 99.7 | 99.7 | 99 |
| JXGC-a | Target | C | 100 | 100 | 99 | 100 | 100 | 96 | 100 | 96 | 100 | 96 | 96 | 100 |
| | | I | 99.8 | 99.9 | 100 | 99.7 | 100 | 100 | 100 | 100 | 100 | 100 | 100 | 99.9 |
| | Query | C | 100 | 72 | 76 | 33 | 100 | 87 | 100 | 87 | 100 | 87 | 87 | 22 |
| | | I | 99.8 | 99.9 | 100 | 99.7 | 100 | 100 | 100 | 100 | 100 | 100 | 100 | 99.9 |
| TX1712-a | Target | C | 87 | 100 | 99 | 100 | 87 | 100 | 87 | 100 | 87 | 100 | 100 | 100 |
| | | I | 99.7 | 99.9 | 99.9 | 99.7 | 100 | 100 | 100 | 100 | 100 | 100 | 100 | 99.9 |
| | Query | C | 96 | 81 | 85 | 36 | 96 | 100 | 96 | 100 | 96 | 100 | 100 | 24 |
| | | I | 99.7 | 99.9 | 99.9 | 99.7 | 100 | 100 | 100 | 100 | 100 | 100 | 100 | 99.9 |
| YCPsy-a | Target | C | 100 | 100 | 99 | 100 | 100 | 96 | 100 | 96 | 100 | 96 | 96 | 100 |
| | | I | 99.8 | 99.9 | 100 | 99.7 | 100 | 100 | 100 | 100 | 100 | 100 | 100 | 99.9 |
| | Query | C | 100 | 72 | 76 | 33 | 100 | 87 | 100 | 87 | 100 | 87 | 87 | 22 |
| | | I | 99.8 | 99.9 | 100 | 99.7 | 100 | 100 | 100 | 100 | 100 | 100 | 100 | 99.9 |
| YNJS7C-a | Target | C | 87 | 100 | 99 | 100 | 87 | 100 | 87 | 100 | 87 | 100 | 100 | 100 |
| | | I | 99.8 | 99.9 | 99.9 | 99.7 | 100 | 100 | 100 | 100 | 100 | 100 | 100 | 99.9 |
| | Query | C | 96 | 81 | 85 | 36 | 96 | 100 | 96 | 100 | 96 | 100 | 100 | 24 |
| | | I | 99.8 | 99.9 | 99.9 | 99.7 | 100 | 100 | 100 | 100 | 100 | 100 | 100 | 99.9 |
| gxpsy-a | Target | C | 100 | 100 | 99 | 100 | 100 | 96 | 100 | 96 | 100 | 96 | 96 | 100 |
| | | I | 99.8 | 99.9 | 99.9 | 99.7 | 100 | 100 | 100 | 100 | 100 | 100 | 99.9 | 99.9 |
| | Query | C | 100 | 72 | 76 | 33 | 100 | 87 | 100 | 87 | 100 | 87 | 87 | 22 |
| | | I | 99.8 | 99.9 | 99.9 | 99.7 | 100 | 100 | 100 | 100 | 100 | 100 | 99.9 | 99.9 |
| Ishi-1-a | Target | C | 87 | 100 | 99 | 100 | 87 | 100 | 87 | 100 | 87 | 100 | 100 | 100 |
| | | I | 99.8 | 99.9 | 99.9 | 99.7 | 100 | 100 | 100 | 100 | 100 | 100 | 100 | 99.9 |
| | Query | C | 96 | 81 | 85 | 36 | 96 | 100 | 96 | 100 | 96 | 100 | 100 | 24 |
| | | I | 99.8 | 99.9 | 99.9 | 99.7 | 100 | 100 | 100 | 100 | 100 | 100 | 100 | 99.9 |
| psy62-c | Target | C | 87 | 100 | 99 | 100 | 87 | 100 | 87 | 100 | 87 | 100 | 100 | 100 |
| | | I | 99.7 | 99.9 | 99.9 | 99.7 | 100 | 100 | 100 | 100 | 99.9 | 100 | 100 | 99.9 |
| | Query | C | 96 | 81 | 85 | 36 | 96 | 100 | 96 | 100 | 96 | 100 | 100 | 24 |
| | | I | 99.7 | 99.9 | 99.9 | 99.7 | 100 | 100 | 100 | 100 | 99.9 | 100 | 100 | 99.9 |
| SGCA5-a | Target | C | 22 | 30 | 28 | 22 | 22 | 24 | 22 | 24 | 22 | 24 | 24 | 100 |
| | | I | 99.8 | 99.9 | 99.9 | 99 | 99.9 | 99.9 | 99.9 | 99.9 | 99.9 | 99.9 | 99.9 | 100 |
| | Query | C | 100 | 100 | 100 | 34 | 100 | 100 | 100 | 100 | 100 | 100 | 100 | 100 |
| | | I | 99.8 | 99.9 | 99.9 | 99 | 99.9 | 99.9 | 99.9 | 99.9 | 99.9 | 99.9 | 99.9 | 100 |

Suppl. Table. 1: **Resemblance of new Type 4 sequences.** The 12 predicted sequences that did not resemble any of the representative phages of type 1, 2 and 3 were compared between them. Putative prophage sequences AHCA1-a and FL17-a were cleaved and the fragment resembling the other sequences was used. Identity (I) and coverage (C) values of two-way local alignments are presented on this table. Green, yellow, and red cells stand for high, medium and low values, respectively (see methods). All sequences were classified as new prophage-like sequence Type 4.

| ORF | Begin | End | UniRef ID | Predicted Function |
|---|---|---|---|---|
| ORF1 | 1 | 480 | WP_015452967.1 | hypothetical protein |
| ORF2 | 483 | 1445 | ONI58489.1 | hypothetical protein |
| ORF3 | 1481 | 1732 | | |
| ORF4 | 1729 | 2073 | | |
| ORF5 | 2075 | 2662 | WP_015452969.1 | hypothetical protein |
| ORF6 | 2672 | 3208 | WP_015452970.1 | hypothetical protein |
| ORF7 | 3209 | 3382 | WP_034441680.1 | hypothetical protein |
| ORF8 | 3396 | 3530 | | |
| ORF9 | 3544 | 4032 | WP_076969211.1 | head protein |
| ORF10 | 4029 | 4538 | | |
| ORF11 | 4713 | 5303 | WP_015452973.1 | hypothetical protein |
| ORF13 | 5554 | 5802 | WP_015452974.1 | hypothetical protein |
| ORF14 | 5932 | 6039 | WP_076969212.1 | phage portal protein |
| ORF15 | 6030 | 6260 | | |
| ORF16 | 6270 | 7322 | | |
| ORF17 | 7399 | 7587 | WP_083965928.1 | DNA packaging protein |
| ORF18 | 7572 | 8327 | | |
| ORF19 | 8314 | 8463 | | |
| ORF20 | 8512 | 8766 | | |
| ORF21 | 8717 | 9085 | WP_015452976.1 | hypothetical protein |
| ORF22 | 10638 | 10889 | WP_015452977.1 | XRE family transcriptional regulator |
| ORF23 | 11723 | 11866 | WP_047263802.1 | DNA helicase |
| ORF24 | 11856 | 12656 | | |
| ORF25 | 12966 | 13112 | OMH86553.1 | head protein |
| ORF26 | 13147 | 13263 | WP_047263803.1 | DUF1376 domain-containing protein |
| ORF27 | 13256 | 13714 | | |
| ORF28 | 14155 | 14247 | AGH16429.1 | ribonucleotide-diphosphate reductase subunit beta |
| ORF29 | 14546 | 14773 | WP_041862253.1 | hypothetical protein |
| ORF30 | 14798 | 14995 | WP_015452983.1 | hypothetical protein |
| ORF31 | 15052 | 15192 | ACT57628.1 | hypothetical protein |
| ORF32 | 15592 | 16056 | WP_015452985.1 | hypothetical protein |
| ORF33 | 17434 | 19329 | WP_015452987.1 | peptidyl-prolyl cis-trans isomerase |
| ORF34 | 20953 | 21690 | WP_015452989.1 | 16S rRNA (uracil(1498)-N(3))-methyltransferase |
| ORF35 | 21761 | 23134 | WP_015452990.1 | glutamate–cysteine ligase |
| ORF36 | 23168 | 23668 | WP_015452991.1 | N-acetyltransferase |
| ORF37 | 23944 | 24588 | WP_015452992.1 | hypothetical protein |
| ORF38 | 25614 | 25793 | ACT57637.1 | hypothetical protein |
| ORF39 | 25806 | 26411 | WP_015452994.1 | pyridoxamine 5'-phosphate oxidase |
| ORF40 | 26554 | 27408 | WP_015452995.1 | enoyl-ACP reductase |
| ORF41 | 27704 | 28756 | WP_015452996.1 | tRNA dihydrouridine(20/20a) synthase DusA |
| ORF42 | 29086 | 30471 | WP_015452997.1 | PTS ascorbate transporter subunit IIC |
| ORF43 | 30511 | 30633 | WP_031935092.1 | hypothetical protein |
| ORF44 | 30713 | 32164 | WP_015452998.1 | deoxyribodipyrimidine photo-lyase |
| ORF45 | 32429 | 33691 | WP_015452999.1 | dicarboxylate/amino acid:cation symporter |
| ORF46 | 33785 | 35980 | WP_015453000.1 | NAD-dependent DNA ligase LigA |
| ORF47 | 36098 | 37765 | WP_015453001.1 | DNA repair protein RecN |
| ORF48 | 37827 | 38642 | WP_015453002.1 | outer membrane protein assembly factor BamD |
| ORF49 | 38769 | 39659 | WP_015824991.1 | UDP-3-O-acyl-N-acetylglucosamine deacetylase |
| ORF50 | 39749 | 41257 | WP_015453004.1 | cell division protein FtsZ |
| ORF51 | 41360 | 42682 | WP_015453005.1 | cell division protein FtsA |
| ORF52 | 42682 | 43596 | WP_015453006.1 | cell division protein |
| ORF53 | 43584 | 44519 | WP_015453007.1 | D-alanine–D-alanine ligase |
| ORF54 | 45122 | 45337 | WP_015453008.1 | hypothetical protein |
| ORF55 | 45453 | 45710 | WP_015453009.1 | hypothetical protein |
| ORF57 | 46336 | 46620 | WP_015453010.1 | DUF4145 domain-containing protein |
| ORF58 | 46765 | 46977 | WP_015453011.1 | hypothetical protein |
| ORF59 | 47803 | 48039 | WP_015453012.1 | hypothetical protein |

| | | | | |
|---|---|---|---|---|
| ORF60 | 48721 | 48849 | ACT57658.1 | hypothetical protein |
| ORF61 | 49539 | 50642 | WP_015453015.1 | terminase |
| ORF62 | 51030 | 51587 | WP_015453016.1 | hypothetical protein |
| ORF63 | 53255 | 54016 | WP_015453017.1 | hypothetical protein |
| ORF64 | 53973 | 54674 | WP_015453018.1 | hypothetical protein |
| ORF65 | 54700 | 55197 | WP_015453019.1 | DUF2800 domain-containing protein |
| ORF66 | 55202 | 55798 | WP_015453020.1 | DUF2815 domain-containing protein |
| ORF67 | 55801 | 57828 | WP_015824993.1 | DNA polymerase |
| ORF68 | 57825 | 58121 | WP_015824994.1 | nuclease |
| ORF69 | 58112 | 59494 | WP_040055333.1 | ATP-dependent helicase |
| ORF70 | 59487 | 59846 | WP_012778351.1 | DNA ligase |
| ORF71 | 59848 | 60420 | WP_015453025.1 | guanylate kinase |
| ORF72 | 60492 | 60800 | YP_007011137.1 | hypothetical protein |

Suppl. Table. 2: **AHCA1-a genes function.** Predicted ORFs for AHCA1-a were annotated using BLASTp against the non-redundant database. Information includes gene start and end position, UniRef ID and gene predicted function.

| ORF | Begin | End | UniRef ID | Predicted Function |
|---|---|---|---|---|
| ORF1 | 1 | 312 | WP_015452960.1 | hypothetical protein |
| ORF2 | 602 | 1738 | WP_015452961.1 | hypothetical protein |
| ORF3 | 1876 | 2328 | WP_045490404.1 | hypothetical protein |
| ORF4 | 2332 | 3048 | KIH96222.1 | hypothetical protein |
| ORF5 | 3303 | 3782 | WP_015452967.1 | hypothetical protein |
| ORF6 | 3785 | 4807 | ONI58489.1 | hypothetical protein |
| ORF7 | 4783 | 5034 | | |
| ORF8 | 5031 | 5375 | | |
| ORF9 | 5377 | 5964 | WP_015452969.1 | hypothetical protein |
| ORF10 | 5974 | 6510 | WP_015452970.1 | hypothetical protein |
| ORF11 | 6511 | 6708 | WP_015824980.1 | hypothetical protein |
| ORF12 | 6846 | 7334 | WP_076969211.1 | head protein |
| ORF13 | 7331 | 7972 | | |
| ORF14 | 8015 | 8605 | WP_015452973.1 | hypothetical protein |
| ORF15 | 9234 | 9341 | WP_076969212.1 | phage portal protein |
| ORF16 | 9332 | 9562 | | |
| ORF17 | 9572 | 10624 | | |
| ORF18 | 10874 | 11629 | WP_083965928.1 | DNA packaging protein |
| ORF19 | 11616 | 11948 | | |
| ORF20 | 12019 | 12387 | WP_015452976.1 | hypothetical protein |
| ORF22 | 13941 | 14192 | WP_015452977.1 | XRE family transcriptional regulator |
| ORF23 | 15159 | 15959 | WP_076969216.1 | DNA helicase |
| ORF24 | 16355 | 16456 | | |
| ORF25 | 16450 | 16566 | WP_047263803.1 | DUF1376 domain-containing protein |
| ORF26 | 16559 | 17017 | | |

Suppl. Table. 3: **Ishi1-a genes function.** Predicted ORFs for Ishi1-a were annotated using BLASTp against the non-redundant database. Information includes gene start and end position, UniRef ID and gene predicted function.

.
